# Performance of Information Theory Derived Semantic Similarity Algorithms for Differential Diagnosis and Clustering

**DOI:** 10.1101/2025.11.17.688933

**Authors:** Ben Coleman, Daniel Danis, Justin Reese, Peter Robinson

## Abstract

Semantic similarity analysis with Human Phenotype Ontology (HPO) enables fuzzy, specificity weighted comparisons of clinical manifestations of individuals and diseases and can be used to support differential diagnostics or to stratify cohorts. Many methods have been proposed to calculate semantic similarity for various applications, including the Phenomizer, which calculates the average best match over all terms in the query and disease, and set-based methods ranging from the Jaccard Intersection to methods that leverage the conditional information content to calculate similarity. However, these methods have not been described under a single mathematical model or robustly compared using a comprehensive data set. Here, we describe several semantic similarity algorithms using derivations based on information theory, propose three of our own variations to these models, and compare the performance of each approach for differential diagnostic ranking and phenotypic clustering.

We find that Phenomizer performs better when diseases are ranked by similarity alone, without generating p-values. Additionally, non-normalized algorithms that use conditional information perform similarly to Phenomizer for differential diagnosis. In contrast, normalized algorithms perform best when clustering cohorts.

**Availability:** Data is available through the Phenopacket-Store (https://github.com/monarch-initiative/phenopacket-store). Algorithms are implemented in the Python package SetSim (https://github.com/P2GX/setsim).

## 1 INTRODUCTION

The advent of precision medicine and individualized medical care has increased interest in precision or deep phenotyping, defined as the precise and comprehensive analysis of phenotypic abnormalities in which the individual components of the phenotype are observed and described [11]. The Human Phenotype Ontology (HPO) has become the *de facto* standard ontology for deep phenotyping in human genetics. The HPO standardizes definitions of abnormal human phenotypes and organizes them using a structured hierarchy that models clinical features as nodes in a directed acyclic graph (DAG). Terms are arranged using IS-A relationships such that each term IS-A subclass of its parent [12, 2].

The HPO provides the foundation for semantic similarity algorithms that quantify phenotypic similarity between a patient and a computational disease model for differential diagnostic decision support as well as for clustering cohorts of patients on the basis of pairwise similarities [7, 9]. Since the publication of the first such method, Phenomizer, numerous other approaches have been presented that leverage information theory in different ways.

Here, we present an information theory-based derivation of set-based semantic similarity functions. We apply these methods to quantify phenotypic similarity and derive our own efficient methods for leveraging conditional information to calculating phenotypic similarity. Finally we compare these methods to the Phenomizer with respect to their performance in differential diagnostic support and clustering. Our methods are freely available as a Python library called SetSim.

## 2. METHODS

### 2.1 Data

#### 2.1.1 Human Phenotype Ontology (HPO) and HPO annotations (HPOA)

Version 2024-12-12 of the hp.json file was used. The HPO project provides computational disease models for currently 8171 Mendelian diseases represented by Online Mendelian Inheritance in Man (OMIM) identifiers. This data was obtained from the phenotype.hpoa file (version 2024-12-12).

#### 2.1.2 GA4GH phenopackets

The GA4GH Phenopacket Schema is a standard for sharing disease and phenotype information characterizing an individual person or biosample that addresses the challenge of documenting case-level phenotypic information [5]. Each phenopacket contains information about the clinical data of one individual including the diagnosis and *T*_*i*_, the set of HPO terms annotated to an individual. We used phenopackets representing 7552 individuals with 481 different diseases; the phenopackets were obtained from release 0.1.22 of the Phenopacket Store repository at https://github.com/monarch-initiative/phenopacket-store [1]. Only phenotype information was used (variant information available in the phenopackets was not used as a part of this analysis)

### 2.2 Information content

Claude Shannon first realized that statistical definitions of entropy could be applied to quantify information. He did this by replacing probabilities of particle states with probabilities of receiving a message, ultimately benefiting from the robust mathematics developed for thermodynamics [13]. For computational deep phenotyping, the information content (IC) of an HPO term is defined based on the frequency of annotation to the term among all considered disease models. Suppose *D* represents the set of all annotated diseases, and *D*_*t*_ the subset of those diseases that are annotated to HPO term *t*. Then, the IC of HPO term *t* being present in a disease or individual (represented by *I* (*t*)) is calculated with the following:

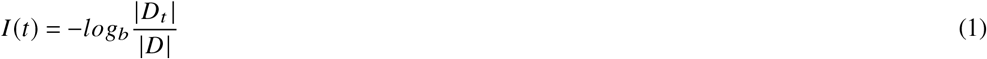

IC is calculated within a given context. Uses of HPO for differential diagnostics calculate IC as given above. It is also possible to calculate the IC of HPO terms based on the distribution of annotations with a cohort. If we represent the cohort as C and individuals within the cohort that are annotated to HPO term *t* as C_*t*_, then the IC can be calculated as

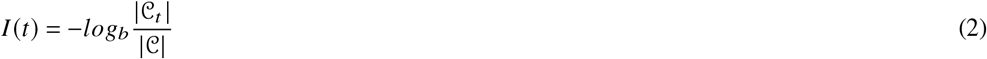

For this work, unless otherwise specified, the IC of each HPO term was calculated according to equation (1) with respect to the entire cohort, with the number of diseases being |*D* | =8324.

### 2.3 True-path rule

The true path rule, also known as the annotation propagation rule, states that an individual who is annotated to some HPO term *t* is implicitly annotated to all of the ancestors of *t* represented by the set of terms *𝒜* (*t*) = *𝒜*_*t*_. For instance, if an individual is annotated to *Zonular cataract* (HP:0010920), that individual implicitly is also annotated to *Cataract* (HP:0000518) and all of the ancestors of cataract. Therefore, a phenopacket representing an individual only needs to include the most specific HPO term while the full set of terms describing that individual, *T*_1_, includes the terms explicitly stated and all ancestors of those terms.

### 2.4 Phenomizer

Phenomizer, the first HPO-based differential diagnosis support algorithm, introduced the concept of using IC to weight clinical features, and to use the hierarchical structure of the HPO to implement fuzzy matching. Phenomizer uses Resnik similarity [10], defining similarity between any two terms as the IC of their most specific common ancestor. For instance, the best match between *Perimembranous ventricular septal defect* (HP:0011682) and *Muscular ventricular septal defect* (HP:0011623) is at their common parent *Ventricular septal defect* (HP:0001629), and their similarity would be calculated as the IC of *Ventricular septal defect*. On the other hand, the best match between *Perimembranous ventricular septal defect* and the unrelated finding *Arachnodactyly* (HP:0001166) would be the root term, *Phenotypic abnormality* (HP:0000118), whose IC is zero. In general, we represent the common ancestor of terms *t*_1_ and *t*_2_ with the highest information content as the most informative common ancestor (MICA), MICA (*t*_1_, *t*_2_).

To quantify the similarity between two sets of terms (either two diseases or individuals), Phenomizer uses a term-by-term approach. The Resnik similarity is found for each term in the first set, *T*_*i*_ (which we represent as the query set *Q* in the following equation), and its most similar term in the second set, *T*_*j*_ (which we represent as the set of terms for disease *D* in the following). The similarity between the two sets is taken as the average of these Resnik similarities.

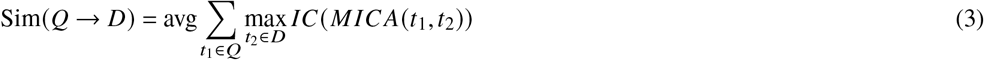

This equation is not necessarily symmetrical, and for some purposes such as clustering, the average of the two directions is taken, Sim (*T*_*i*_,*T*_*j*_) = 0.5 · Sim (*T*_*i*_ →*T*_*j*_) + 0.5 · Sim (*T*_*j*_ →*T*_*i*_).

The raw similarity score depends on a number of factors, including the number and specificity of the terms both of the query and of the diseases represented in the database. When first introduced, it was shown that the performance of this model could be improved by calculating an empirical p-value for the observed similarity score, corresponding to the probability of obtaining a given similarity score or better by choosing the same number of query terms at random [7].

### 2.5 Set-based Phenotype-driven Semantic similarity methods

An alternative to Phenomizer’s term-by-term approach which treats each term annotation in a set independently, it is possible to consider all terms in each set simultaneously to calculate a similarity between sets. The set-based methods can be divided into four classes depending on whether similarity is calculated as an unweighted intersection of terms, or if the terms are weighted by their IC. Further, these methods can be divided according to whether the intersection is normalized by the union of terms (Table 1). In the following sections, we present a consistent mathematical notation for specifying each of these algorithms. We leverage the insight gained from the comparison to implement novel algorithms that we call SetSim and NSetSim.

**Table 1.**
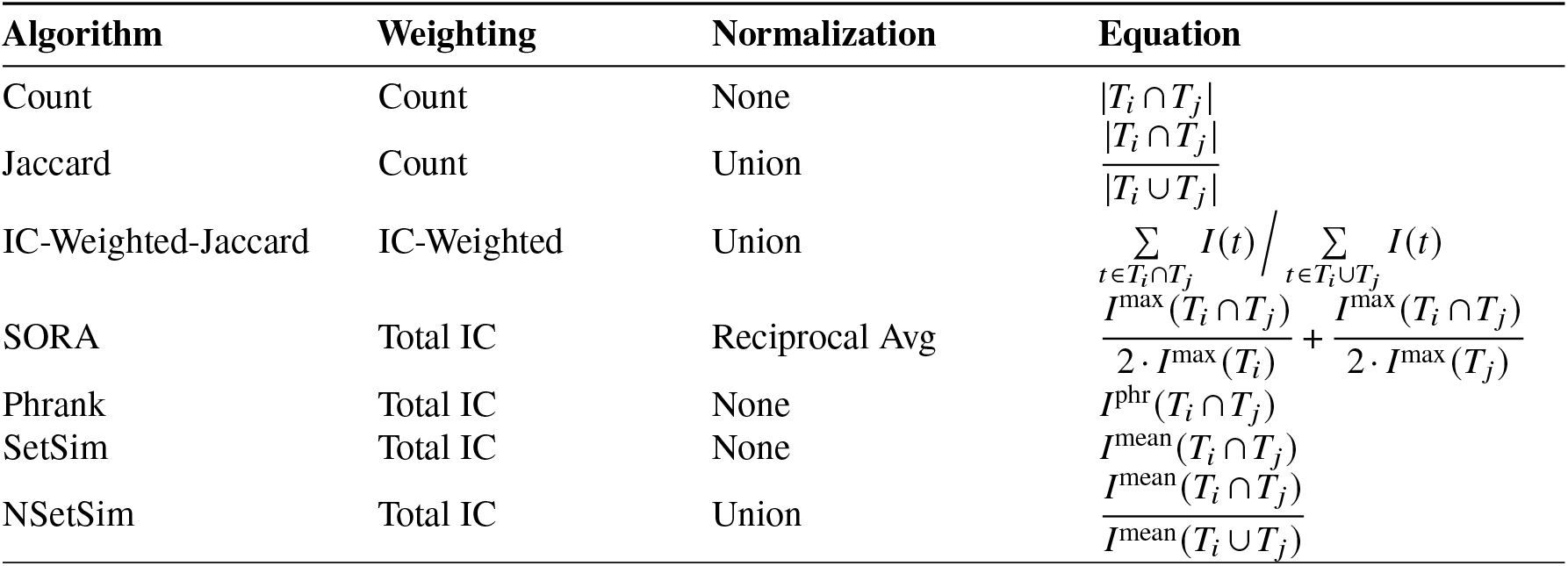
Summary of Set-based Semantic Similarity Methods for Phenotype-driven Differential Diagnosis.

#### 2.5.1 Intersection count

The simplest possible way to calculate similarity would be to count the number of terms in the intersection set. We can model this with the following where *T*_*i*_ and *T*_*j*_ represent the full sets:

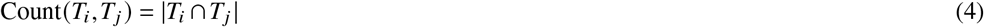

#### 2.5.2 Jaccard index

The similarity defined by equation (4) can be normalized using the Jaccard method by dividing by the cardinality of the union [4].

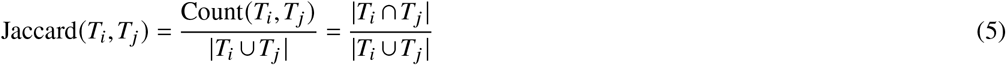

The Jaccard index has been used to calculate gene function similarity in the algorithm SimUI [3].

#### 2.5.3 IC-Weighted-Jaccard

The algorithm SimGIC [8] adapts the basic Jaccard index by weighting each term by its IC using the following, where *I* (*t*) is the IC of the term *t*:

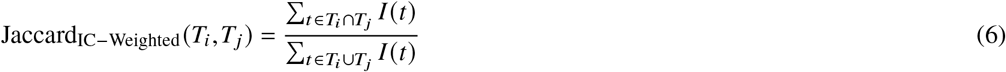

For simplicity we will refer to this approach as IC-Weighted-Jaccard. Weighting terms in this way can be used to account for inconsistencies in ontological depth and the differences between adjacent nodes in a graph.

### 2.6 Calculating the Information of Multiple Terms

The information of several events (the presence of several terms in this case) can be calculated by summing the information of each event if the events are independent. In the case of the HPO, the terms are arranged in a hierarchical IS-A structure and are not independent because of the true-path rule. To determine the total information of the intersection or union sets, we therefore use the following equation where *t*_1_, *t*_2_, …, *t*_*i*_ is the set of terms, *T*. For conciseness, we will use the shorthand *I* (*t*) to imply *I* (*t* ∈ *T*):

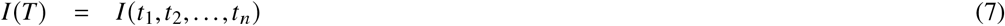

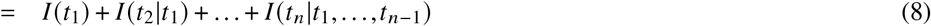

To solve this equation, we must solve for the conditional information using the IS-A relationships encoded in our ontology. For instance, if *t*_*a*_ is an ancestor of *t*_*i*_, then the information of *t*_*a*_ given *t*_*i*_ is zero, i.e., *I* (*t*_*a*_|*t*_*i*_) = 0, because we already know that *t*_*a*_ is present given that *t*_*i*_ is present. By extension, if *t*_*i*_-_1_, …, *t*_1_ are all ancestors of *t*_*i*_, then the information of the set is equal to the information of *t*_*i*_. For a more complex set of terms, we utilize the following equation where *t* _*p*_ is the only direct parent of *t* and *𝒜* _*t*_ represents the set of ancestors of *t*_*i*_:

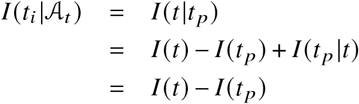

We can extend this equation for multiple parents where *𝒫* _*t*_ (*t*) = *𝒫* _*t*_ is the set of parents of *t*:

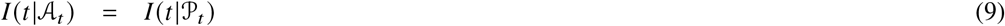

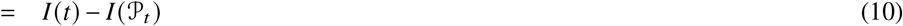

Using the above, we can easily sum the conditional information of each term starting with the information of the root term and iteratively take the information of each descendant minus the information of its immediate parent.

#### 2.6.1 Calculating similarities between sets of HPO terms

Because of the true-path rule, an annotation to an HPO term implicitly includes all of its ancestors. Therefore, if a query, a representation of an individual patient, or a disease model is annotated to a set of specific HPO terms, then the set also includes all ancestors of that set. The following four methods (SORA, Phrank, SetSim, and NSetSim) calculate the similarity between two sets of HPO terms (*T*_*i*_ and *T*_*j*_) is based on the information in the intersection of the terms.

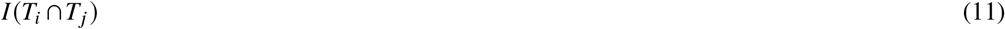

For differential diagnostic queries, *T*_*i*_ represents the HPO terms observed in an individual and *T*_*j*_ represents the HPO terms in a disease model. For clustering, *T*_*i*_ and *T*_*j*_ represent the HPO terms observed in individuals *i* and *j*.

The four methods differ in how they calculate this quantity for terms which have multiple parents, using either the maximum, mean, or a Bayesian approach, as explained in the following text.

#### 2.6.2 SORA

The approach described in section 2.6 was first introduced in the algorithm SORA [14]. SORA solves conditional probabilities in the reverse direction by starting with leaf terms (terms with no children in a set). The authors introduced the concept of extended IC:

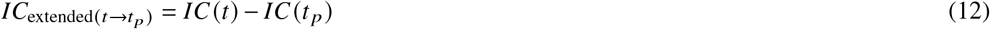

SORA solves the challenge of calculating this quantity starting iteratively from the leaves of the graph. This is simplified by the fact that any set with only one leaf can be solved by finding the IC of that one term. The advantage of this approach is that the IC of a term is always equal to the total IC of the term and its ancestors.

A complication when looking at HPO terms with two or more parents. For example, the dual parentage of *Arachnodactyly*. We can model the case of dual parentage with the equation below where *t* _*p*1_ and *t* _*p*2_ represent the parents of *t*_*i*_:

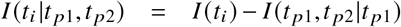

Notice that if the IC of *t*_*i*_ is less than the total IC of the parents, then the resultant conditional information will be negative. This occurs because this approach assumes orthogonality between *I* (*t* _*p*1_ | *t* _*p*2_) and *I* (*t* _*p*2_ |*t* _*p*1_) but HPO does not guarantee this. This approach is computationally demanding compared to other approaches we will describe, but we can approximate this method by taking the maximum IC of the parents rather then the total IC. This will occasionally result in slightly higher conditional ICs but removes the possibility of negative conditional ICs. We will describe two additional algorithms to calculate total IC that are mathematically equivalent to this except in the case of dual parentage.

SORA is unique from other approaches in that it uses a reciprocal average to normalize similarities. This can be represented with the following:

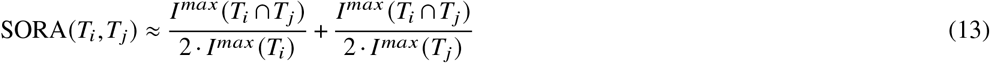

In this notation, *I* (*T*_*i*_ ∩ *T*_*j*_) refers to the information content of the intersection between the sets of terms *T*_*i*_ and *T*_*j*_ (e.g., between a query and a disease model or between two individuals). The IC of each term in the intersection is calculated as described above.

#### 2.6.3 Phrank

An alternative for finding the conditional information of terms with multiple parents that avoid negative conditional information is used in the algorithm Phrank [6]. Except for the case of multiple parents, Phrank is mathematically equivalent to SORA.

Still, instead of solving for conditional information in the information space, Phrank calculates the conditional information of each term given its parents by finding the IC of the probability of a term given its parents. This is done using the following equation where *D*_*t*_ is the set of diseases annotated with *t* and *D* _*p*_ is the set of diseases annotated with at least one parent:

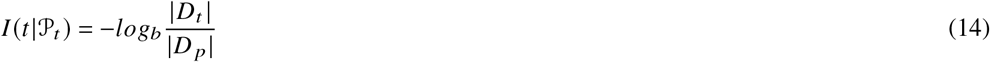

Of the methods described in this manuscript, Phrank will always produce the highest conditional information of terms with multiple parents. This is because Phrank calculates the conditional information a term given any parent rather than all parents, which is more intuitive given that all parents are present by the true path rule. This results in the total information of a term and its ancestors being greater than its IC if that term or its ancestors have multiple parents. Additionally, Phrank is uniquely restricted because |*D*_*p*_| cannot be determined from information content in the case of multiple parentage. This is because we need to know the set of diseases that are shared among parents so they are only counted once.

The Phrank procedures uses the above definition to calculate the total information content in of all terms in the intersection of the query and the disease model.

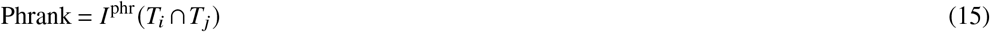

#### 2.6.4 SetSim

Here we present a novel approach that we call SetSim for calculating the conditional information of a term as the IC of that term minus the mean IC of the parents. This will reduce the inflation of a term’s IC seen when using Phrank and prevent negative conditional information.

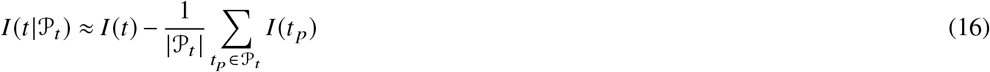

In summary, we calculate the similarity between sets of terms and their ancestors by finding the intersecting set of terms and calculating the total information content of the set of terms, which is equal to the sum of the conditional information of each term given its parents.

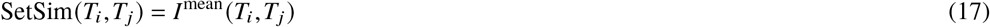

#### 2.6.5 NSetSim

In analogy to Jaccard normalization of the counting method, we can normalize SetSim by dividing by the total information of the union of sets. We will call this NSetSim and describe it using the following:

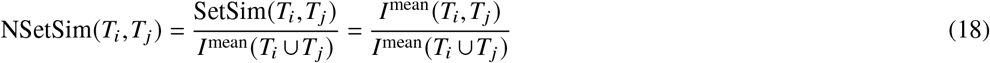

### 2.7 P-Value Generated Rankings

Unlike other semantic similarity methods we have discussed, Phenomizer is commonly used with an additional layer for ranking diagnoses to a set of phenotypic features. This is done by randomly selecting a large number (e.g., 100,000) of sets of phenotypic features and finding the similarity of each set to a disease. An individual’s similarity to a disease will then be compared to distribution of the random sets of terms in order to calculate an empirical p-value. The Phenomizer algorithm uses similarities to break ties for diseases assigned the same p-value. To account for differences in similarity from the number of features in the set, Phenomizer determines p-values using 100,000 sets of terms with the same number of terms as the set of interest [7]. In principle, this method can be used for any of the algorithms above.

### 2.8 Testing Diagnostic Ranking

To test these algorithms, we created SetSim, a Python library available under a BSD-3 Clause License, to conveniently find the semantic similarity of individuals whose phenotypic features are encoded as phenopackets. Using SetSim we compare Phenomizer with the four types of intersection-based algorithms described, as well as the two new approached SetSim and NSetSim.

Because of the computational demand of calculating p-values for every possible number of terms, we limited the samples to those with 30 or fewer terms. Individuals were ranked according to all diseases, and the rank of the correct diagnosis was returned. The average rank for each method was plotted and statistical significance comparing the performance of each algorithm was done using Mann-Whitney U test with SciPy (version 1.10.1) [15].

To compare how similarity measures would perform on individuals with many irrelevant features, we identified the set of common HPO terms that were present in at least 10% of diseases (*IC* < *log* (10)). We then added a random selection of 20 of these terms to each patient. Redundant terms were removed. We repeated the analysis using the samples with added noise. Additional samples were removed when filtering for samples with 30 or fewer terms, resulting in a set of 7550 samples using original phenopackets and 7060 samples when adding noise. (Adding terms for noise resulted in some samples crossing the 30 term threshold.)

### 2.9 Clustering with Pairwise Similarity

We additionally measured the performance of each algorithm for identifying clusters of similar patients. We prepared sets of patients from the phenopacket-store with similar diseases. First, we identified diseases with at least 30 diagnosed patients in our sample, resulting in 65 sets of patients. From these, we randomly selected 5 sets for each clustering experiment.

We re-calculated the IC and conditional IC of terms after removing the five selected diseases from the hpoa file (simulating the case where patients share an undiscovered diagnosis). This restricted us to using a common method for calculating conditional IC for terms with multiple parents. As a result, Phrank was removed from this analysis, and SORA was replaced with RASetSim, a version of NSetSim that uses reciprocal average for normalization similar to SORA. Then, using each similarity algorithm, we followed the methods proposed by Reese et al. for clustering patients with Phenomizer [9]. First, we created a pairwise similarity matrix for the individuals using each similarity algorithm. Treating each row as a vector representing a patient, we clustered the patients using k-Means clustering with five clusters for the five diseases. To score the clustering, we labeled each cluster with one of the 5 diseases based on the prevalence of individuals with that disease. We then calculated the total number of correctly clustered patients whose diagnosis corresponded to their cluster. We report the fraction of patients who were correctly clustered using each strategy. To investigate this for the case of patients with many common terms, we repeated the analysis using the same samples but with 20 common terms added using the methods described above. Using a bootstrap method, we repeated this process 1,000 times iterating through different sets of 5 randomly selected diseases.

### 2.10 Software availability

SetSim is available under an MIT license at https://github.com/P2GX/setsim. A tutorial and example notebook are provided.

## 3 RESULTS

In this work, we compare eight methods for calculating semantic similarity in the context of phenotype-driven differential diagnosis (Table 1). We provide an information theory-based discussion of commonly used methods and a comparison of the performance of the methods on differential diagnostic support and clustering.

### 3.1 Performance of semantic similarity methods for differential diagnosis

We compared the performance of several algorithms for finding semantic similarity using phenopackets representing 7552 individuals with 481 diseases. For each individual, the rank of the correct diagnosis was recorded, and the distribution of ranks for all individuals was compared between methods.

Using original samples (without artificial noise), Phenomizer performed significantly better than all algorithims except SetSim and Phrank. Interestingly, count outperformed methods that used a normalization step despite not incorporating IC. When adding artificial noise, SetSim and Phrank outperform Phenomizer. The counting method performs more similarly to normalized approaches when artificial noise is added. In general, normalized approaches performed significantly worse than non-normalized approaches. SORA (which uses a reciprocal average normalization) performed between normalized and non-normalized approaches. There was no difference in performance of SetSim compared to Phrank suggesting that variations of conditional IC for terms with multiple parents has little effect on performance (Figure 1).

**Figure 1.**
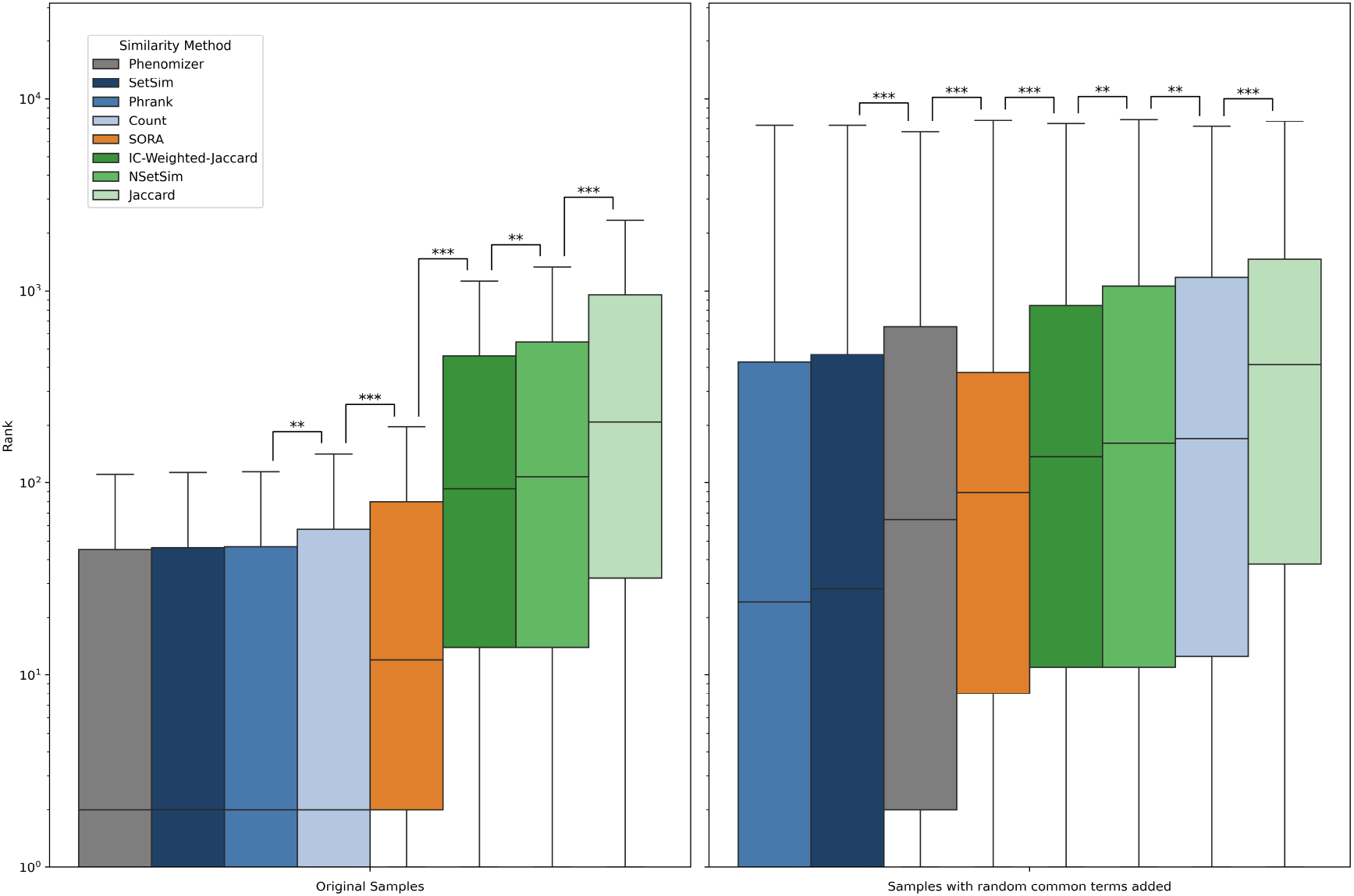
Rank Comparison of Similarity Algorithms. Box plots show the distribution of ranks for the correct diagnosis for 7552individuals (lower numbers represent better performance). Each algorithm is with original phenotypes and phenotypes with 20 common terms added for noise. Methods are ordered by Mann-Whitney U-statistic with asterisks indicating a significant difference. Colors are organized by normalization method (green for union, orange for reciprocal average, and blue for none) with the lighter hues representing count based methods.

The original Phenomizer algorithm uses empirical p-values to rank diseases for a patient. We compared the performance of Phenomizer ranking by p-value vs ranking by similarity alone. There is a significant improvement in performance when similarity is used alone, without ranking by empirical p-values. This result may be related to the substantially increased HPO annotation results available currently compared to the resources available when Phenomizer was published in 2009.

### 3.2 Performance of semantic similarity methods for clustering cohorts

We compare the performance of seven similarity measures for clustering individuals by their pairwise similarity (Figure 3). For software performance, all methods calculated conditional information for terms with multiple parents by subtracting the mean IC of those parents. As a result we represent Phrank and SetSim together and SORA as RASetSim, a version of NSetSim that normalizes with reciprocal average. For original samples without added noise, RASetSim (approximately SORA) outperformed other algorithms. For samples with random terms added to simulate noise, all normalized methods performed similarly to Phenomizer and outperformed non-normalized methods. (Figure 3).

**Figure 2.**
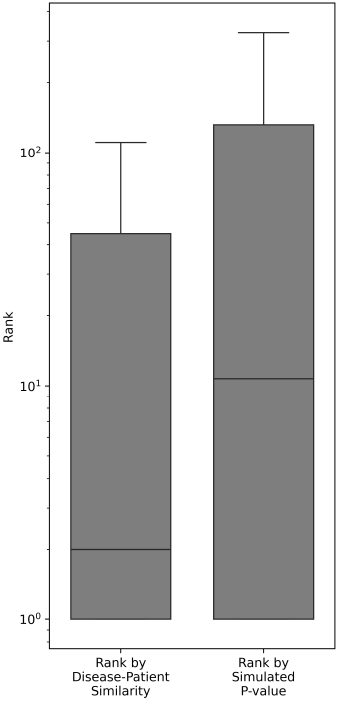
Phenomizer using Similarity vs Simulated P-value Rankings. Box plots show performance of Phenomizer using the original algorithm which uses simulated p-values to rank diseases vs a simpler method that uses disease-patient similarity directly. Results show that Phenomizer’s performance is improved when disease-patient similarity is used directly without using simulated p-values.

**Figure 3.**
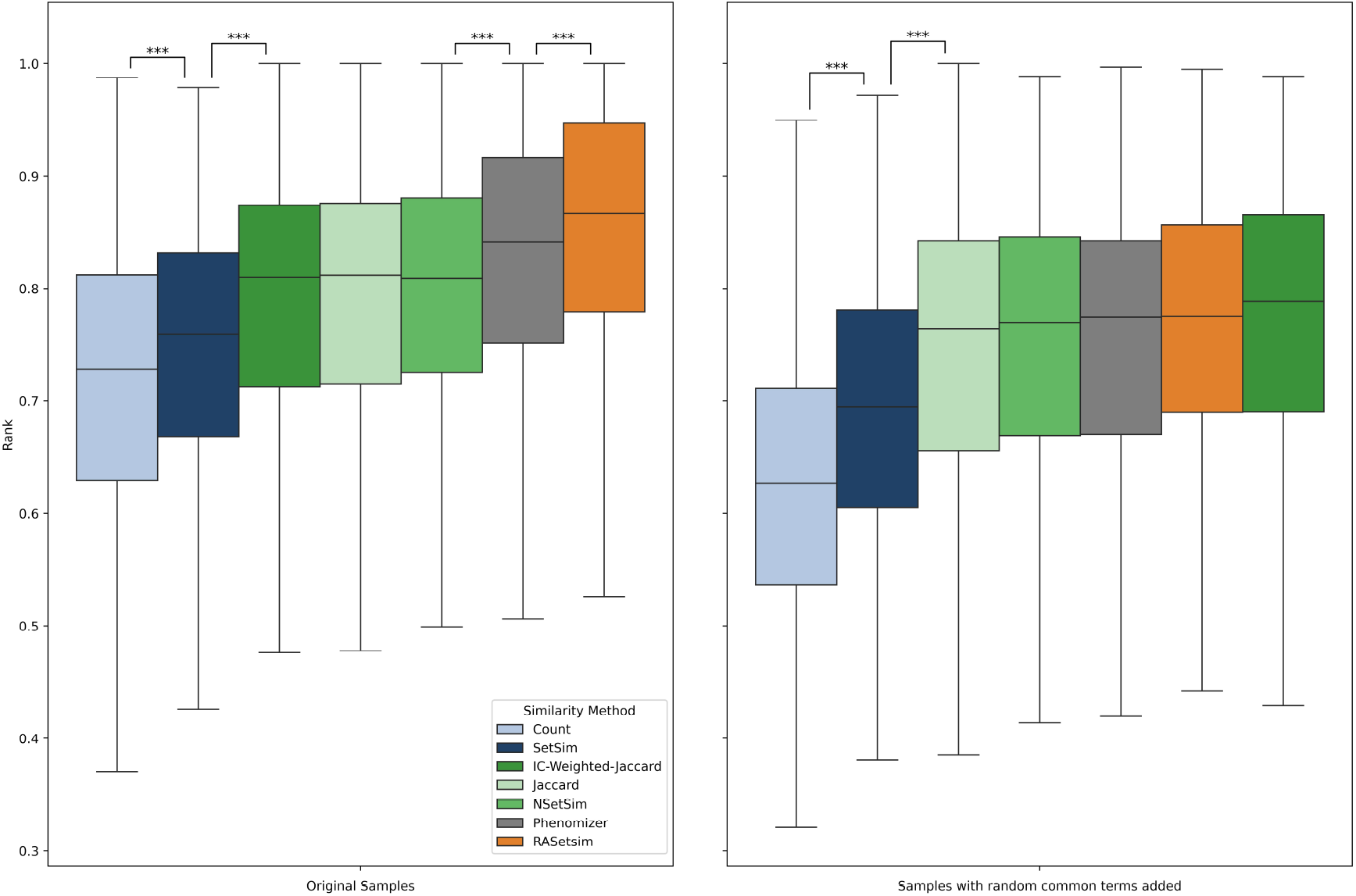
Performance of Person-to-Person Clustering. We compare the performance of K-Means clustering using a pairwise similarity matrix generated from each algorithm shown. Clustering scores represent the fraction of patients correctly clustered according to diagnosis averaged from 1,000 bootstrap iterations. Clustering was tested on original phenotypes and phenotypes with 20 common terms added for noise. Methods are ordered by Mann-Whitney U-statistic with asterisks indicating a significant difference. Colors are organized by normalization method (green for union, orange for reciprocal average, and blue for none) with the lighter hues representing count based methods.

## 4 DISCUSSION

Semantic similarity is the basis for many algorithms the leverage HPO for clustering or differential diagnostic support. Numerous algorithms and software packages have been published since the initial publication of the Phenomizer in 2009. The methods for these algorithms were presented in disparate ways, making it difficult to understand the similarities and differences between the various methods. In this work, we have compared Phenomizer against seven other approaches, including four set-based approaches and three straightforward approaches (count, Jaccard, IC-weighted Jaccard). We present a common mathematical framework for understanding the methods. Perhaps unsurprisingly, no one method performs best in all situations. Here, we provide a discussion of the factors we consider most important for determining the performance of a semantic similarity algorithm.

### 4.1 The relative influence of common and specific HPO terms

Phenomizer averages similarities across terms (See equation 3). Therefore, the highest similarities are achieved when we only include the most relevant and informative term. Including common terms in a patient’s phenotype (even if they are not symptoms of the patient’s disease) may decrease the calculated similarity. Alternatively, intersection-based approaches (which use the total of shared phenotypes) will make similarity increase or stay the same when more terms are added to an individual’s phenotypic profile. This makes set-based similarity approaches more tolerant to comparing patients with many common terms, such as may be the case when extracting HPO terms fromsources like electronic health records (EHRs) which include many nonspecific phenotypic features. This may be related to the observation that the *p*-value approach that displayed superior performance in 2009 is no longer superior to the raw-score approach used with the original Phenomizer algorithm [7].

### 4.2 Weighted and unweighted intersection approaches

We can expect that weighted approaches will offer an improvement over their unweighted counterparts because they give greater similarity between sets where the intersection contains fewer common terms. Our results show that weighting becomes more important when samples include many unrelated common terms. Interestingly, weighting secondary to normalization method when looking at differential diagnosis with original samples. To our surprise, IC-weighted-Jaccard (SimGIC) outperformed the more nuanced approaches such as NSetSim. In future studies, it would be interesting to explore variations of IC-weighted-Jaccard that use alternative methods of normalization. The more nuanced differences of calculating conditional IC of terms with multiple parents does not lead to significant differences, suggesting that the more flexible, information theory-based approaches we apply in SetSim are superior to alternatives.

### 4.3 Differential diagnostics versus clustering

Our results show that non-normalized approaches perform better for differential diagnoses, whereas normalized approaches perform better for clustering. The former is likely a reflection of the comprehensive nature of HPO annotations. More conservative annotations would include phenotypic features that are highly associated with the disease and therefore have a higher negative predictive value (NPV) when absent. However, HPO annotations include less common features of a disease which have a low NPV. Normalized methods will penalize the absence of these uncommon features despite them having low NPV, whereas non-normalized methods will ignore them.

By this same logic we can see why normalized methods work better when clustering. Within a cluster of individuals with a shared disorder, you can observe the frequency of features of that disorder. Clustering with normalized similarity methods will invoke a greater penalty for the absences features with higher NPV (features shared among most individuals in a cluster) and a smaller penalty for features with low NPV.

### 4.4 Phenomizer

Phenomizer performs worse when ranking based on p-value. This is interesting given that when Phenomizer was created, it was shown to perform better using p-value. This could be due to a number of factors. The first is that we used published clinical descriptions (case reports or cohorts with detailed clinical information) for this test, while Phenomizer was originally tested using simulated data. It is also possible that the 8,000 new terms added to the HPO since the creation of Phenomizer have made random samples of terms less likely to be similar to a specific disease. As a result, null distributions will become less effective for discriminating between individuals that are similar to diseases. An additional consideration is the importance of p-values for significance testing for diagnostics. Rather than ranking with p-values, diseases could be ranked with similarity, and a multiple testing corrected p-value could be produced based on the highest-ranked disease.

When we use Phenomizer without calculating p-values, we find that it is the only method that performs in the top three methods for all applications tested. While specific applications may benefit from certain methods, Phenomizer remains the state-of-the-art for general use.

### 4.5 Conclusion

No single semantic similarity algorithm is superior for all applications. The performance of the algorithms is likely to depend on the structure of both computational disease annotations and the depth of patient phenotyping. Published algorithms make use of concepts from information theory in related ways. For diagnostic tasks, Phenomizer performed best for data sets with no added noise. Phrank and our SetSim approach showed superior performance with random common terms added with the intention of modeling noise as might be found in EHR data. For clustering, in contrast, normalized approaches and Phenomizer performed best. Our work provides important insights for choosing algorithms and approaches for phenotype-driven differential diagnostic support and clustering. We have implemented all algorithms in an open-source Python package called SetSim that is freely available at https://github.com/P2GX/setsim.

## ACKNOWLEDGMENTS

The presentation of theory and algorithms given here was refined through discussion following a research progress talk with the Department of Genetics and Developmental Biology at UConn Health.

